# Experimental evolution of hybrid populations to identify Dobzhansky-Muller incompatibility loci

**DOI:** 10.1101/2023.05.24.542153

**Authors:** Nicole Szabo, Asher D. Cutter

## Abstract

Epistatic interactions between loci that reduce fitness in inter-species hybrids, Dobzhansky-Muller incompatibilities (DMIs), contribute genetically to the inviability and infertility within hybrid populations. It remains a challenge, however, to identify the loci that contribute to DMIs as causes of reproductive isolation between species. Here, we assess through forward simulation the power of evolve and resequence experimental evolution of hybrid populations to map DMI loci. We document conditions under which such a mapping strategy may be most feasible and demonstrate how mapping power is sensitive to biologically relevant parameters such as one-way versus two-way incompatibility type, selection strength, recombination rate, and dominance interactions. We also assess the influence of parameters under direct control of an experimenter, including duration of experimental evolution and number of replicate populations. We conclude that an evolve and re-sequence strategy for mapping DMI loci, and other cases of epistasis, can be a viable option under some circumstances for study systems with short generation times like *Caenorhabditis* nematodes.

## Introduction

A major challenge to understanding the process of speciation is to determine the genetic basis to how natural selection, genetic drift, recombination, and other factors interact to generate and maintain distinct biological entities. The process of speciation starts when subsets of individuals from the same species become reproductively isolated from each other, often as a by-product of geographic barriers (Savolainen et al. 2013, Stankowski and Ravinet 2021). The limitation to gene flow that ensues allows genetically intrinsic mechanisms of reproductive isolation to evolve, namely pre-zygotic barriers, which prevent mating or fertilization, and post-zygotic barriers, which act after mating by yielding infertile or inviable hybrid offspring (Presgraves 2010a, Maheshwari and Barbash 2011, Stankowski and Ravinet 2021). Hybrid infertility and inviability caused by genetically intrinsic factors can permanently restrict the exchange of genes between populations to make hybrid individuals evolutionary “dead-ends” that then serves to enforce reproductive isolation and the evolutionary formation and maintenance of distinct biological species (Coughlan and Matute 2020). Dobzhansky-Muller incompatibilities (DMIs) are now appreciated as an important genetic basis for the evolution of such intrinsic post-zygotic isolation in speciation and yet the characterization of their location and identity in genomes remains challenging (Coyne and Orr 1998a, Presgraves 2010b, Castillo and Barbash 2017).

More than 80 years after Dobzhansky’s pioneering work, the genes responsible for reproductive isolation remain incompletely characterized across organisms (Castillo and Barbash 2017). However, recent technological developments and the increasing accessibility of genome-wide sequencing approaches have allowed for the isolation of specific DMI genes in laboratory settings (Maheshwari and Barbash 2011, Schumer et al. 2014, Blackman 2016, Powell et al. 2020). Such approaches include the introgression methods first pioneered by Dobzhansky, genome-wide association studies (GWAS), quantitative trait locus (QTL) mapping, clinal analysis, and linkage disequilibrium (LD) analyses (Barton and Hewitt 1989, Payseur 2010, Gompert and Buerkle 2011, Fitzpatrick 2013, De La Torre et al. 2015). LD mapping, in particular, has proved especially useful in determining not only the location but the number of loci participating in incompatible interactions (Schumer and Brandvain 2016, Schumer et al. 2020), as demonstrated in swordtail fish by quantifying hundreds of DMI pairs in the naturally occurring hybrid zones of two species (Schumer et al. 2014, Schumer et al. 2015, Powell et al. 2020). Specifically, they exploited patterns of correlation between allelic ancestry and linkage in the hybrid genomes in what is termed an ancestry disequilibrium scan: unlinked markers of the same parental origin that demonstrate linkage disequilibrium in hybrid populations indicate the presence of an incompatibility pair.

An analogous experimental approach to ancestry disequilibrium mapping for elucidating DMIs involves evolve and re-sequence (E&R) experiments (Kofler and Schlötterer 2014, Long et al. 2015). E&R, as applied to mapping speciation genes (Pereira et al. 2021), combines multi-generation experimental evolution of hybrid populations with Pool-Seq, a modified next-generation high-throughput DNA sequencing approach that economically allows for the sequencing from a population pool of many individuals (Turner et al. 2010, Kofler et al. 2011, Turner et al. 2011, Ferretti et al. 2013, Kofler and Schlötterer 2014). While hypotheses on DMIs have rarely been tested directly in E&R studies, E&R has been applied to explore speciation and epistatic genetic interactions more generally (White et al. 2020). For instance, E&R using *Escherichia coli* to characterize the type of epistasis at play permitted the conclusion that negative epistasis contributes to intraspecific variation (Maharjan and Ferenci 2013). The potential for E&R to provide a means to successfully map inter-species DMIs across genomes, however, remains to be fully explored.

Given the substantial resource and time investments required to carry out E&R, simulations provide a cost-effective means to guide long-term wet-lab experiments that aim to characterize DMI loci (Schumer et al. 2020). One such simulation study assessed the power to map DMIs in yeast populations using linkage analysis, finding that large sample sizes and certain statistical methods performed best in successfully detecting DMI loci (Li et al. 2013). More recently, simulations explored the likelihood of hybrid speciation by elucidating how dominance, recombination, and selection influence DMI resolution (meaning that one or both alleles involved in an incompatibility pair get purged), though the power to localize DMI loci per se was not assessed (Blanckaert and Bank 2018). Simulations of DMI inference from ancestry disequilibrium scans showed how ongoing hybridization of hybrid populations to parental populations can lead to biased co-ancestry in the genome (Schumer and Brandvain 2016).

Here, we conduct forward-time computer simulations to explore the power of experimental evolution to detect and localize loci that contribute to epistatic genetic incompatibilities. By mimicking the *Caenorhabditis* nematode system that has partially-isolated species pairs (Woodruff et al. 2010, Dey et al. 2014, Bundus et al. 2018, Cutter 2018), we ground these simulations in a biological context with the potential to guide future evolve and re-sequence experiments while also assessing general properties of the potential for E&R mapping of DMIs to succeed. To date, *Caenorhabditis* has been used in experimental evolution across tens to hundreds of generations to study the evolution of sex, mutation rates, development, pathogen interactions, population genetic theory, thermal sensitivity, and other traits (Gray and Cutter 2014, Teotónio et al. 2017), but not to map DMIs in hybrid populations. In contrast to co-ancestry disequilibrium scan approaches (Schumer and Brandvain 2016), we aim to detect signatures at individual loci by taking advantage of replicated hybrid populations within an E&R framework. By exploring a range of parameters that define intrinsic properties of a system as well as features manipulable by an experimenter, we identify conditions most amenable to successful mapping of DMI loci with E&R experiments.

## Methods

### Evolving Hybrid Populations in SLiM

To test the power of evolve and re-sequence experimental evolution studies to map DMIs, we simulated the evolution of hybrid populations containing Dobzhansky-Muller incompatibilities (DMIs) using SLiM software (3.3.1) (Haller and Messer 2019). We created artificial autosomal genomes containing 1) a single two-locus DMI or 2) a suite of 10 randomly-placed two-locus DMIs, then tracked their evolution in populations that mimic the hybridization of two species. Each replicate population contained 1000 diploid individuals, which stayed constant across generations. Over time, genotypes that created DMIs were selected against and purged from the populations, leading towards fixation of one or the other parental allele which we then attempted to map from patterns of parallel allele frequency changes across replicated hybrid populations.

Each individual’s genome started heterozygous at each locus, reflecting the logic that F_1_ individuals that initialized the simulations would have inherited a distinguishable allele from each parental species (Figure 1). In the two-locus model, each locus of the DMI locus pair was linked to a separate autosome, with 10,000 evenly-spaced neutrally-evolving loci across the two autosomes as markers. No new mutations entered the populations over time. In the multi-locus model, the 20 loci that comprise 10 DMI pairs were randomly placed along two autosomes among an additional 10,000 evenly-spaced neutrally-evolving loci. Because DMI locus positions in the multi-locus model were randomly assigned, they may overlap with neutral-evolving loci in some populations.

**Figure 1.**
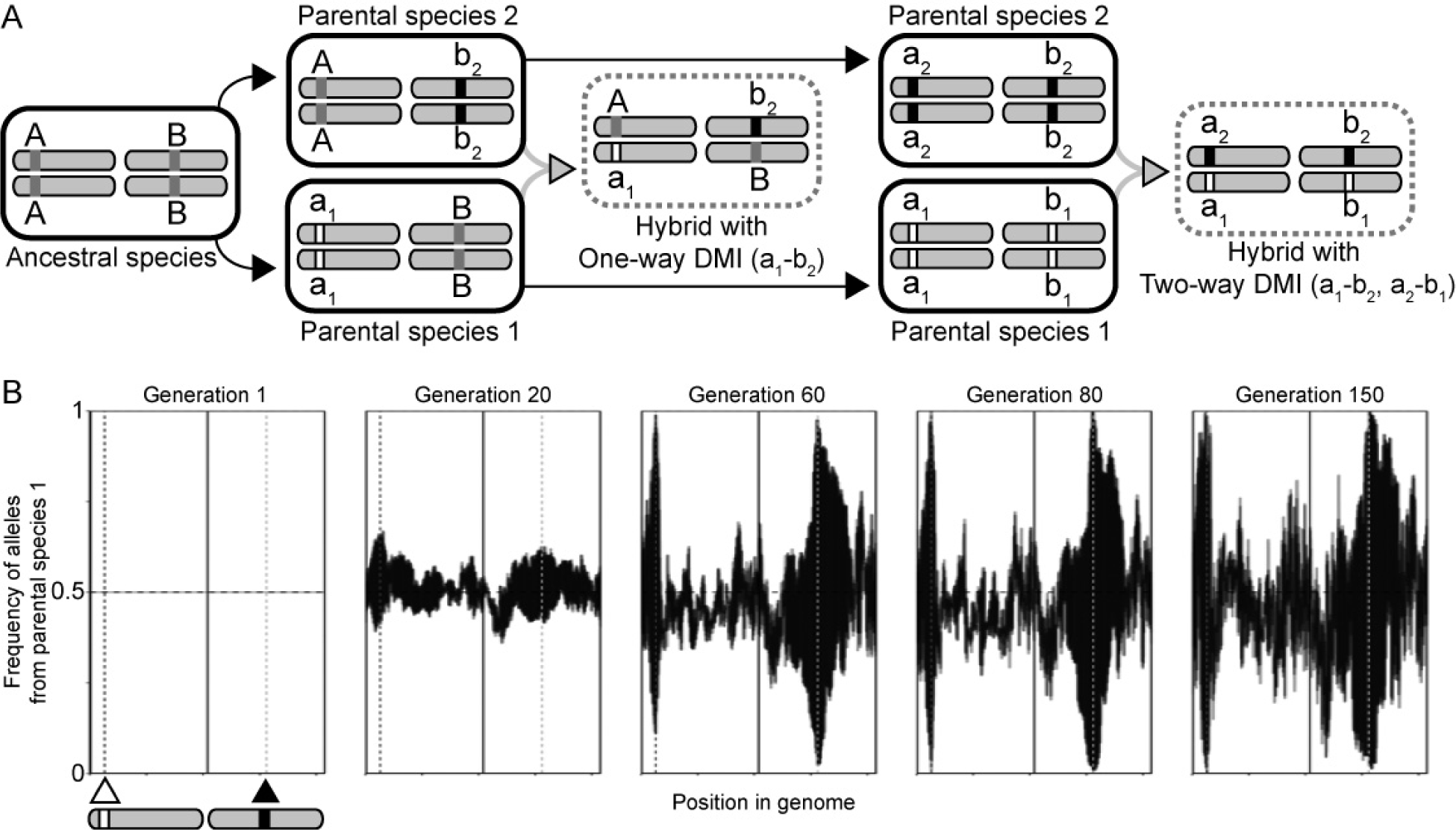
(A) Schematic representation of the evolution of one-way and two-way DMIs from a single ancestral species. Our simulations were initiated with F1 hybrid individuals derived from the two parental species. (B) Simulated frequencies of alleles from parental species 1 over time in five replicate hybrid populations. Triangles and vertical dashed lines indicate positions of loci involved in a two-way DMI. While this example data was created using the same script as the simulated populations containing two-way DMIs in the results, it was generated to demonstrate the methods used and not for analysis. At generation 1, all hybrid individuals in each population are F1 heterozygotes (allele frequency 0.5). In this example of two-way DMI, resolution of the DMI occurs upon fixation of alleles of both loci from parental species 1 or, alternately, upon fixation of alleles of both loci from parent species 2. The superimposed allele frequency profiles for the 5 replicate populations represented thus shown either extreme high or extreme low frequency near the DMI loci.

Each 20.8 Mb chromosome mimics the typical length of a *Caenorhabditis* chromosome (Teterina et al. 2020). Simulated genomes mimic the chromosome architecture of *Caenorhabditis* nematodes with a central region of low recombination and flanking arms of higher recombination (Rockman and Kruglyak 2009). Experimental evolution studies in *Caenorhabditis*, including *C. elegans* and *C. remanei*, commonly are run for tens to hundreds of generations (Gray and Cutter 2014, Teotónio et al. 2017). Previous crosses between *C. latens* males and *C. remanei* females demonstrated X-autosome (causing hybrid male sterility), mito-autosomal, and autosomal-autosomal incompatibilities (the latter two responsible for inviability) (Bundus et al. 2018). The simulations, however, only consider autosome-autosome incompatibilities that are otherwise neutral, meaning that they do not individually confer a fitness advantage over alternative alleles. We also did not simulate explicitly the acquisition or analysis of short read sequence data, however, but assume that the simulated neutral marker loci represent distinguishable variants extracted from an evolve and re-sequence experiment. In this way, the simulation output may be used to guide the design of experiments with organisms such as *C. latens* and *C. remanei* to map hybrid incompatibilities between them.

We traced the evolutionary fate of alleles at DMI loci under a range of epistatic selection coefficients (*s* = 0.05, 0.2, 0.8 across all DMI pairs), interaction types (one-way or two-way interactions for the two-locus scenario), number of DMIs (multi-locus or two-locus scenarios with one-way interactions), and generation time sampling points (10 to 500 generations). Selection coefficients were chosen to allow us to investigate plausibly detectable effects ranging from mild to severe. For simulations of a single DMI pair, we encoded one DMI locus on an arm of one chromosome and the DMI partner locus on the centre of the other chromosome (the high recombination region “HRR” of 10 cM/Mb and low recombination region “LRR” of 0.1 cM/Mb, respectively). For simulations of multiple DMI pairs, the DMI loci were randomly encoded throughout the two autosomes, irrespective of recombination regions; otherwise, the autosomes were constructed identically to those of the two-locus model. Parameter values were inspired by the biological range of what is known in *Caenorhabditis* species about chromosomal recombination landscapes and hybrid fitness (Rockman and Kruglyak 2009, Dey et al. 2014, Bundus et al. 2018, Teterina et al. 2020). The simulated recombination rates are slightly more extreme than empirical averages, allowing these simulations to provide both high and low bounds with respect to less extreme recombination environments.

In modeling one-way incompatibilities, we followed the classical depiction of DMIs that occur when each parental species contributes only one allele to a derived-derived DMI pair (Figure 1). Alternate allele combinations at the two loci would be compatible, because they would include ancestral alleles shared by the two species (Figure 1). In two-way incompatibilities, by contrast, each allele contributed by a parent at a DMI locus is incompatible with both of the alleles that could be contributed by the other parent at the partner DMI locus. Two-way DMIs thus result in two incompatibility types involving four derived alleles at two loci (Figure 1), which can occur when distinct mutations arose and fixed at both loci in each species separately after they diverged from their common ancestor, or from negatively-epistatic ancestral-ancestral and ancestral-derived allelic interactions (Figure 1).

For simplicity in the multi-locus case involving 10 pairwise one-way DMIs, we simulated only one dominance scenario such that both DMI alleles of a pair acted dominantly. For one-way DMIs of the two-locus model, by contrast, we considered four dominance scenarios. First, both DMI alleles at each locus acted dominantly (dominant × dominant), resulting in expression of the incompatibility regardless of whether an individual was homozygous or heterozygous at a given DMI locus, and manifesting in the F_1_ generation. Second, both DMI alleles were recessive (recessive × recessive), resulting in expression of the incompatibility only when an individual was homozygous for both DMI alleles and thus manifesting only in the F_2_ and later generations. For the final two dominance scenarios (dominant × recessive), dominance differed by recombination region: the DMI allele in the high recombination region was recessive and the DMI allele in the low recombination region was dominant, or vice versa. The 10 DMI locus pairs conferred additive fitness effects, such that, for example, in the moderate selection case, each DMI decremented relative fitness by 0.02 for a maximum possible reduction of 20% across all 10 DMIs.

For the two-locus model with two-way DMIs, we simulated just two dominance scenarios. In one scenario, all four DMI alleles at each locus were dominant, resulting in the expression of both negative interactions starting in the first generation. Fitness effects from each incompatibility were multiplicative. For example, in the moderate selection case, for *s* = 0.106 for both DMIs present in an individual, fitness was reduced by 20% (80% of ancestral fitness) versus the 10.6% reduction (89.4% of ancestral fitness) when only one negative interaction was present. In the second scenario, all four DMI alleles were recessive, resulting in the expression of the DMIs only in later generations. Unlike in the first dominance scenario, both negative interactions could not be expressed at the same time in a given individual, resulting in a maximum fitness reduction of 10.6% when *s* = 0.106.

### Estimating DMI Locations

Our simulations mimicked an evolve and resequence experiment by evaluating the power to detect DMIs from parallel allele frequency changes across replicate populations. We considered experimental evolution of 5, 10, and 20 replicate populations used to map DMIs for a given parameter set. Using an R script that processed allele frequencies to assess parallel evolution across a given set of replicate populations, we attempted to infer DMI locations on the simulated autosomes, iterating this procedure for each of 500 independent sets of population replicates to assess the precision and accuracy of mapping estimation (github.com/Cutterlab/Generating-DMI-Estimates) (Figure 1). All simulation runs were carried out via *gnu-parallel* on the University of Toronto’s Niagara/SciNet supercomputer (Ponce et al. 2019).

Estimates of the DMI locus locations were based on allele frequencies that were consistently more extreme than what would be expected from genetic drift alone. In some cases, this procedure inferred no DMI. To infer the location of DMI loci, we searched for overlapping genomic regions with extreme frequency values among the replicate populations. Allele frequencies were deemed extreme in a given replicate population when they exceeded values plausibly caused by genetic drift alone. The drift cut off to define extreme allele frequencies was determined from 2.5th and 97.5th percentiles of 1000 neutral simulations that otherwise resembled the non-neutral simulations, including linkage structure, except that the alleles at the “DMI loci” exerted no fitness impacts. After counting the number of replicate populations with an extreme value at a given locus, we calculated with *stats* package in R the binomial probability of retrieving the observed number of extreme populations for each locus given the number of replicate populations considered (i.e., 5, 10, or 20 replicate populations), flagging loci with probabilities <0.05 after false discovery rate correction.

In the two-locus scenario, genomic positions within the 2-log ∼95% confidence interval of the maximum negative log-binomial p-value for each chromosome were included to define the edges of the DMI location. If no loci significantly demonstrated an extreme number of populations with extreme allele frequencies for one or both chromosomes, then no DMI was estimated for that set of replicate populations (thus contributing to imperfect “detectability”). We then assessed whether the estimated DMI locations were accurate based on whether they overlapped with the correct DMI positions input into the simulations. The precision of the estimates was calculated as the length of the regions that accurately overlapped true DMI loci divided by the length of the chromosome. This process was repeated 500 times for each distinct parameter set to assess the overall variability, detectability, accuracy, and precision of mapping DMI locations in the simulated genomes. We report the accuracy for a parameter set by tallying the number of estimates that overlapped the true DMI loci and dividing by the number of estimates made (usually 500, except when detectability was imperfect) and the precision for the parameter set by calculating the median relative length of the estimated DMI intervals.

In the multi-locus DMI simulations, we marked estimated DMI locus positions as blocks of adjacent loci with flagged probabilities <0.05, provided that they included at least two loci. If two or more blocks of loci occurred within 6240 bp of each other (a span of 2.5 neutral loci), they were merged into a single estimate of DMI locus position. We report the number of loci inferred to be mapped, the incidence of true-positive detection among the estimated DMI mappings, and the width of the regions spanning estimated DMI positions to assess the accuracy and precision of mapping DMI loci in genomes simulated to have 20 pairs of DMI loci.

## Results

### Weak-to-moderate strength one-way DMIs can be mapped feasibly with E&R

When considering a single pair of DMI loci that express one-way incompatibility, the estimated map positions most accurately overlap the true DMI loci for intermediate durations of experimental evolution (Figure 2A). The precision of those map positions, however, increases with the number of generations of experimental evolution (Figure 2B). These general trends hold for weak to strong fitness effects, dominance regimes, and high versus low recombination environments; as expected, mapping precision is greater in high-recombination regions for a given duration of experimental evolution (Figure 2B). The dominance of interactions showed at most a negligible effect on overall mapping accuracy and precision (Figure 2). Mapping accuracy and precision, however, are most sensitive to the duration of experimental evolution when selection is strong (*s* = 0.8), showing best accuracy for durations ≤100 generations and best precision for durations ≥100 generations. With weak selection (*s* = 0.05), by contrast, the relatively flat sensitivity to duration of experimental evolution yields the best trade-off between accuracy and precision between 40 and 200 generations. For both strong and weak selection on DMI loci, the precision of mapping is extremely poor and highly variable if populations experience 40 generations or fewer (the mapping interval often comprises >70% of the length of the chromosome), with the inference method unable to detect any DMI loci at all in 16% of runs after 10 generations with weak selection (Figure S6). Intermediate selection on DMI loci (*s* = 0.2), however, permits robust accuracy and precision of DMI locus mapping from 10 to 100 generations in high recombination regions, with mapping intervals spanning as little as 8% of the chromosome length (Figure 2B). Overall, an E&R approach may be most suitable to mapping pairwise one-way DMI loci of weak-to-moderate effect in a timeframe of 40 to 100 generations of experimental evolution.

**Figure 2.**
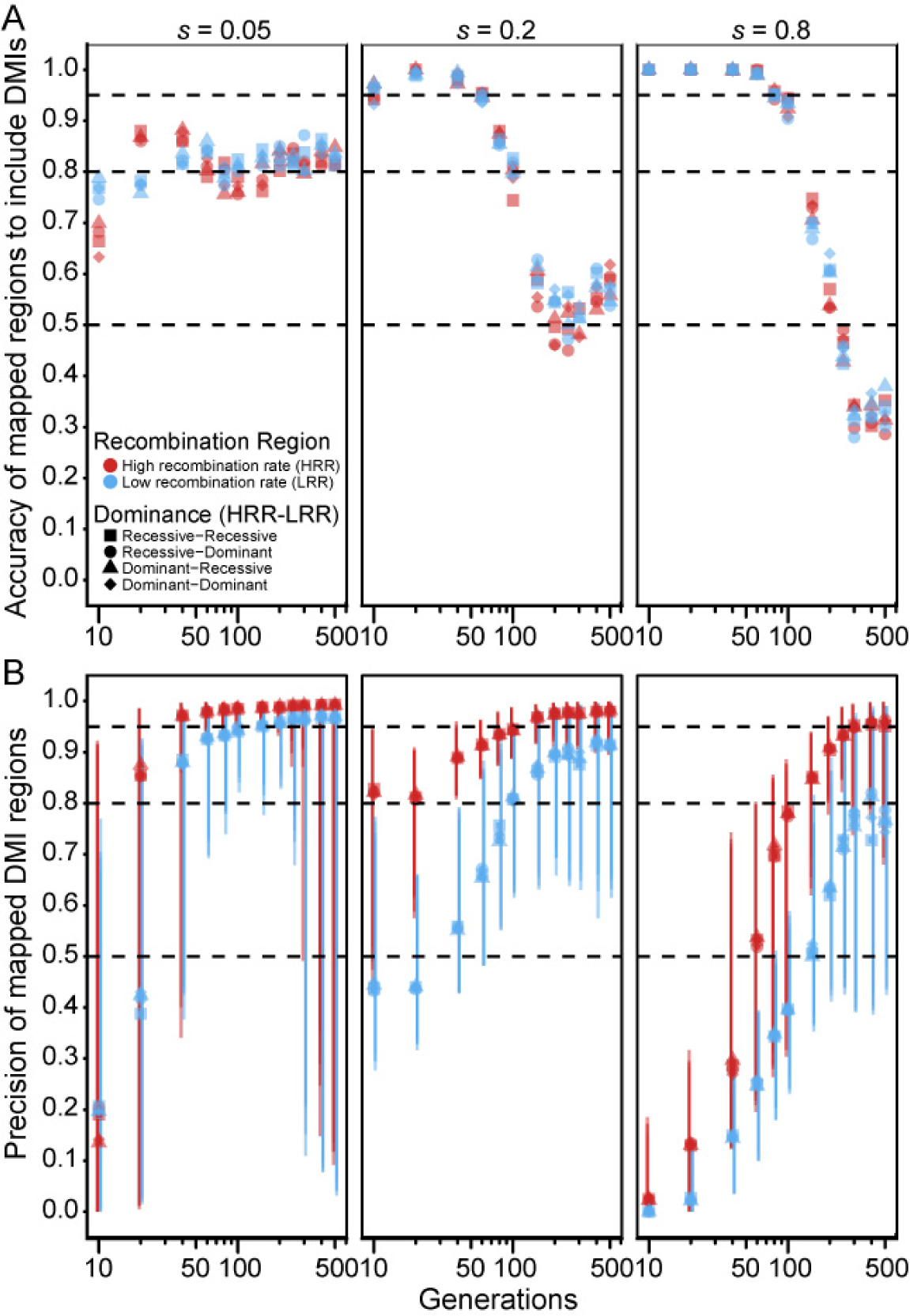
(A) Accuracy and (B) precision of DMI mapping as a function of duration of experimental evolution in generations (log scale) for one-way incompatibilities. High recombination region (HRR, red points) and low recombination region (LRR, blue points) are shown for four different dominance landscapes (symbol shapes). Simulations map DMI loci based on parallel allele frequency changes among 10 replicate populations per parameter set. Accuracy is the number of mapped estimates that overlap true DMI loci divided by the total number of inferred estimates. Precision is the median width (bp) of correct estimates divided by the length of one chromosome. Error bars around the median correspond to the 95% interquantile range (2.5^th^ and 97.5^th^ quantiles of mapped widths). Horizontal dashed lines correspond to accuracy and precision values of 50%, 80%, and 95%.

### Accurate and precise mapping of two-way DMIs with E&R requires 100 generations or more

When considering a single pair of DMI loci that express two-way incompatibilities, such that both alleles at one locus experience reciprocal negative interactions with both alleles at the other locus (Figure 1), we observed that both precision and accuracy of the estimated map positions tend to increase with the number of generations of experimental evolution (Figure 3; Figure 4). This pattern contrasts with the trends for one-way DMI loci that showed a trade-off between accuracy and precision (Figure 3A and 4A cf. Figure 2A). The improving accuracy and precision of two-way DMI mapping with duration of experimental evolution holds for weak to strong fitness effects, dominance regimes, and high versus low recombination environments. Similar to one-way DMI loci, mapping precision is greater in high-recombination regions for dominant DMIs for a given duration of experimental evolution (Figure 3B). For instance, when selection is strong, we observed that mapping precision in high-recombination regions reaches a peak value of 96.5% versus a peak value of 83.1% for low-recombination regions. For recessive DMIs, however, a discernible beneficial influence of recombination on mapping precision holds true only when selection is strong (Figure 4B). More generally, we observe a clear interaction between dominance and strength of selection on the accuracy and precision of the estimated map positions for two-way DMI loci. When selection is moderate or weak on incompatibilities, mapping precision tends to be better for dominant than for recessive DMI loci for any given duration of experimental evolution and, in absolute terms, mapping resolution of recessive incompatibilities is poor with fewer than 150 generations. Strong selection, by contrast, shows greater mapping precision for recessive two-way DMIs for all but the shortest durations of experimental evolution and with only modestly reduced accuracy for durations of 40 generations and less. When recombination is high, we observed that mapping precision reaches a value of 96.1% by generation 100 for recessive DMIs (Figure 4B), compared to 78.4% for dominant DMIs (Figure 3B). Strong selection on two-way recessive DMIs in high recombination regions thus provide the most promising circumstances for mapping two-way DMI loci, with nearly 90% precision and accuracy after just 20 generations of experimental evolution (Figure 4). However, most other circumstances that we explored require 100 generations or longer for high accuracy and precision in mapping two-way DMI loci.

**Figure 3.**
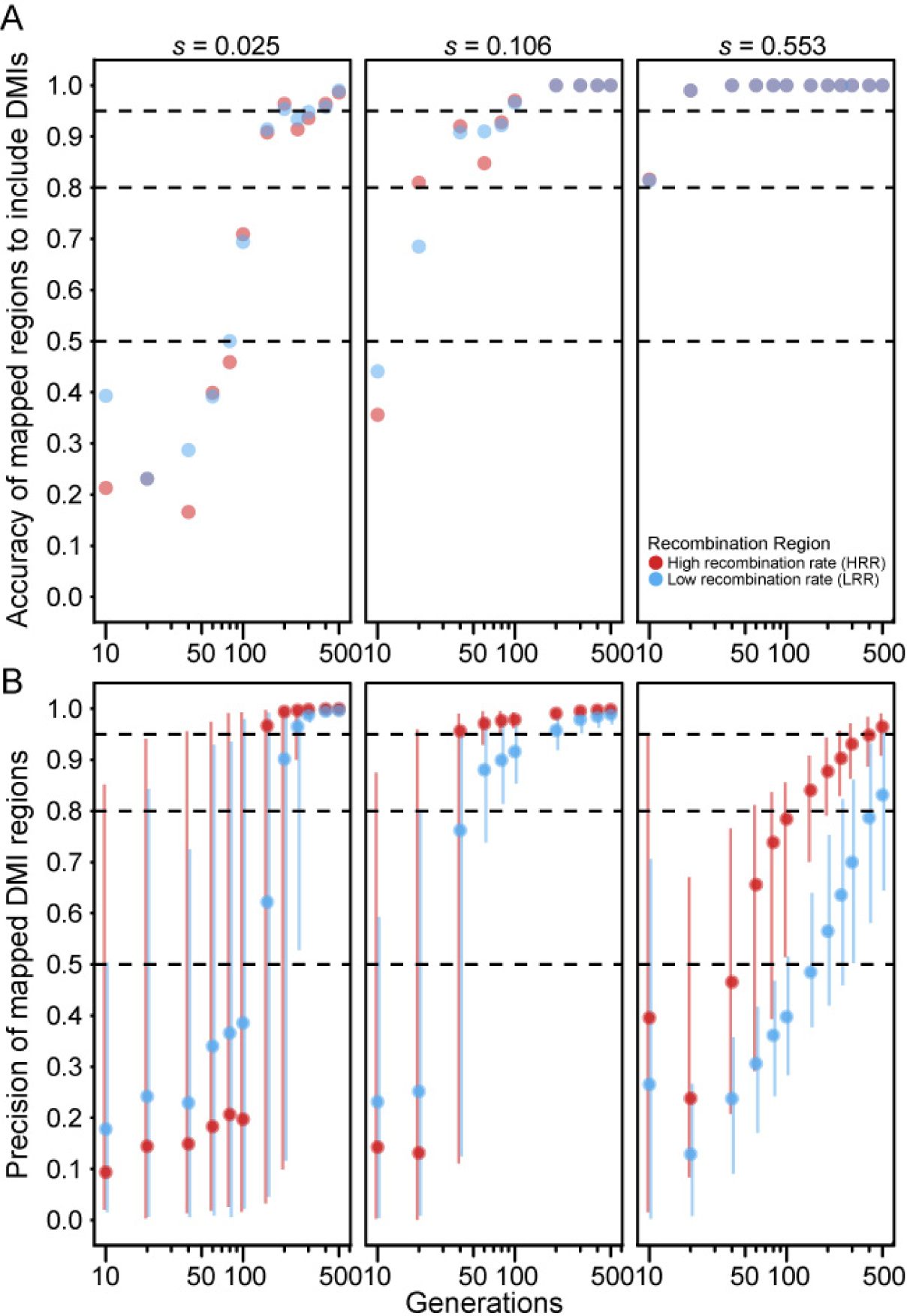
(A) Accuracy and (B) precision values across generations of hybrid experimental evolution (log scale) for dominant × dominant two-way incompatibilities in the high recombination region (red) and low recombination region (blue). Accuracy, precision, error bars, and dashed lines as in Figure 2. Simulated selection parameters (*s*) per DMI are smaller for two-way DMIs relative to one-way DMIs (cf. Figure 2) to account for multiplicative fitness effects to retain a consistent maximum fitness reduction of 0.05, 0.2, or 0.8.

**Figure 4.**
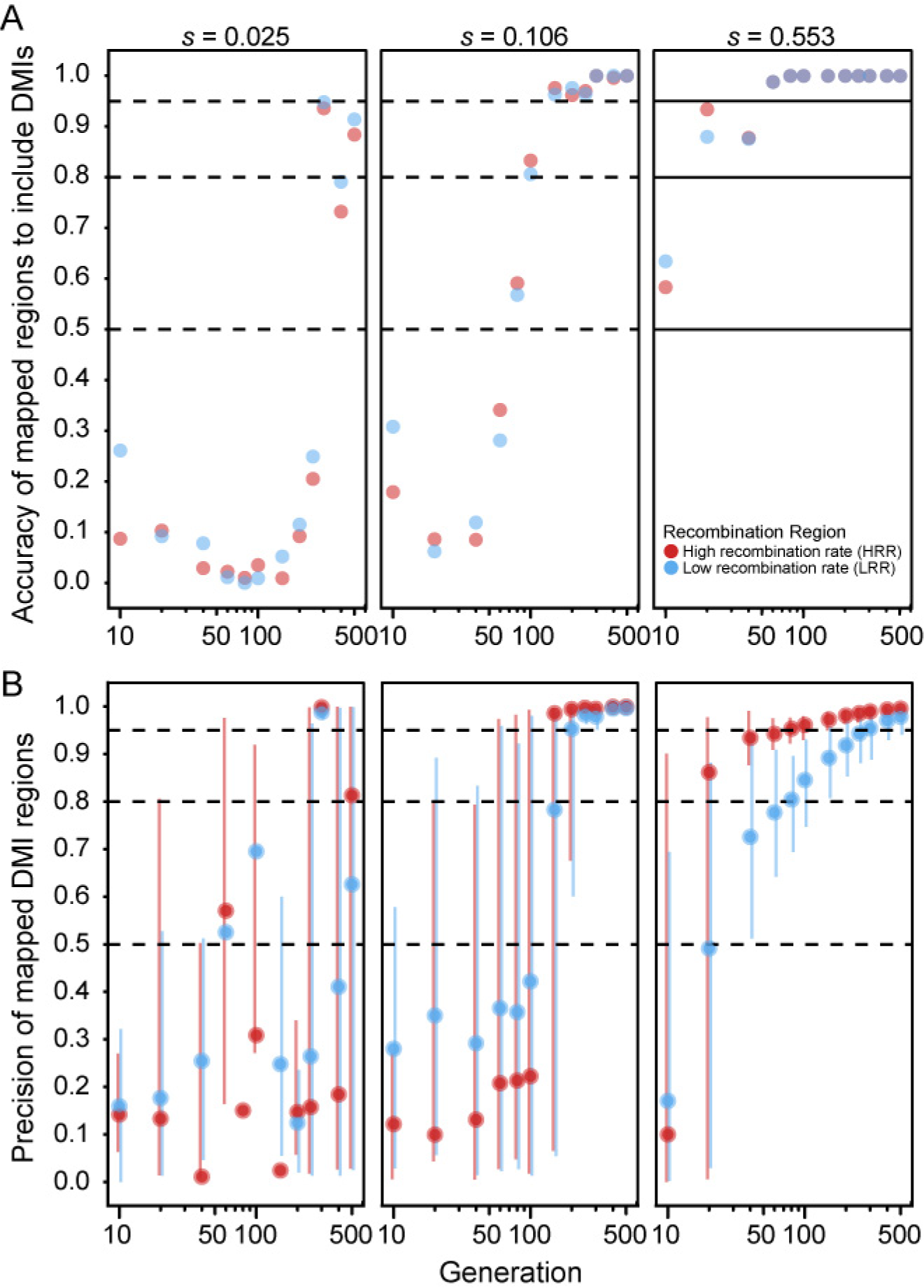
(A) Accuracy and (B) precision values across generations of hybrid experimental evolution (log scale) for recessive × recessive two-way incompatibilities in the high recombination region (red) and low recombination region (blue). Accuracy, precision, error bars, and dashed lines as in Figure 2. Simulated selection parameters (*s*) per DMI are smaller for two-way DMIs relative to one-way DMIs (cf. Figure 2) to account for multiplicative fitness effects to retain a consistent maximum fitness reduction of 0.05, 0.2, or 0.8. There is no low recombination point for the parameter set *s* = 0.025 at 80 generations because there were no accurate estimates retrieved for this parameter set (median accuracy = 0), and precision of mapped DMI regions are only calculated across accurate estimates.

### Complex trade-offs between true- and false-positive mapping with many pairs of DMI loci

Finally, we considered genomes with 20 DMI loci (10 pairs) that express one-way incompatibilities, such that just one allele at each locus experiences reciprocal negative interactions with one allele from another locus. We observed that the median size of estimated map locations tends to decrease with the number of elapsed generations (Figure 5A), consistent with the pattern of increasing precision observed in simulations of just a single pair of one-way DMI loci (Figure 2B). Median estimated size of DMI regions is not very sensitive to the number of replicate populations, though simulations of the fewest replicate populations (five) tend to have widest genomic spans around estimated DMI locations. When selection on each DMI pair is weak (*s* = 0.005) or moderate (*s* = 0.02), mapping intervals are quite narrow (median estimated width does not exceed 0.2% of the genome). When selection is strong (*s* = 0.145), however, mapping resolution is entirely uninformative for the first 40 generations (Figure 5A). Specifically, we observed 100% of the genome typically to be implicated in incompatibility, regardless of the number of population replicates, though eventually reaching high precision after 60 generations or more (achieving mapping resolution for strong selection below 0.1% of chromosome length after 200 generations with 20 replicate populations; Figure 5A; Figure S1).

**Figure 5.**
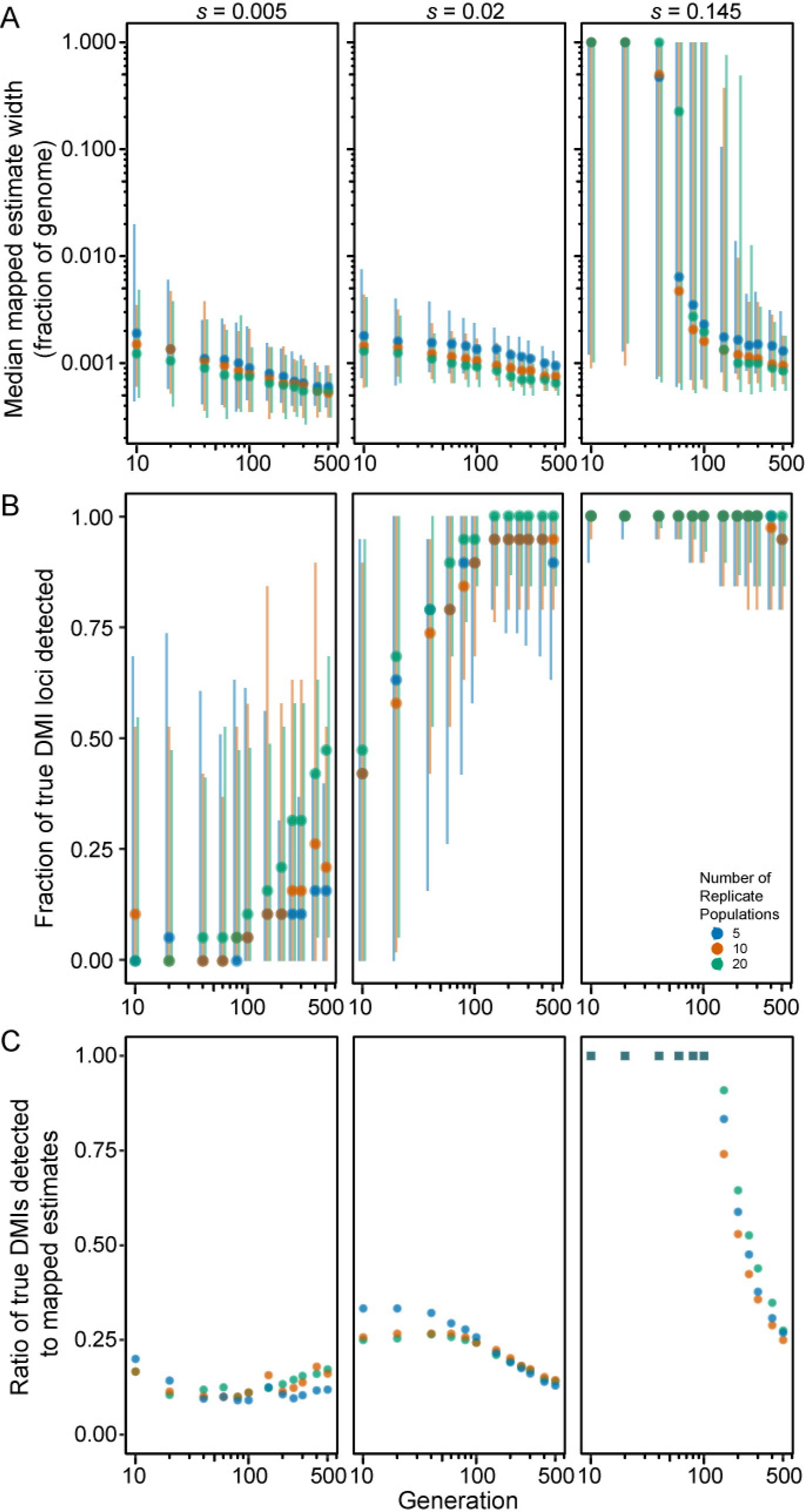
Power to detect DMI loci in experimentally-evolving populations with 10 pairs of one-way DMI loci. (A) Median mapped estimate width (log scale fraction of genome), (B) proportion of true DMI loci detected, and (C) ratio of the number of true DMI loci mapped to the number of mapped regions, as a function of duration of experimental evolution (log scale generations) for 20 DMI loci that express one-way incompatibilities for 5, 10, and 20 replicate populations. Error bars indicate 2.5^th^ to 97.5^th^ quantiles. Square symbols in (C) indicate parameter sets for which mapped regions overlapped >1 true DMI on average.

As the regions spanning estimated DMI locations get narrower as generations elapse, the number of true DMI loci detected also increases (Figure 5B), reaching 90% or more of the 20 DMI loci after 100 generations with moderate selection (*s* = 0.02 for five replicate populations). However, the total number of estimated DMIs also increases with time, surpassing the true number of DMI loci, and leading to an accumulation of false positives (Figure S2). Consequently, we observed that the fraction of true positive estimates of DMI location exhibits a complex relationship with time and selection strength (Figure 5C). When selection is strong, true positive detection of DMIs is highest in earlier generations when the regions span most of the genome and so have poor resolution. In later generations, true positives decline to account for approximately 25% of mappings by generation 500 when the roughly 20 estimated DMI locations each represent about 0.1% of the length of a chromosome (Figure 5C). With strong selection, essentially all 20 DMI loci occur within the estimated regions, irrespective of the number of generations elapsed (Figure 5B). When selection is moderate (*s* = 0.02), the proportion of true positive estimates for DMIs is highest with just 5 replicate populations until generation 150 (Figure 5C), as a result of the fewer estimates made with this lower powered design (Figure S2). However, the rapid growth of DMI estimates over time with moderate selection leads to a decline in true positives down to ∼14% by generation 500, irrespective of the number of population replicates (Figure 5C). As a result, early generations permit detection of only about ∼45% of the 20 true DMI loci, reaching about ∼ 100% of true DMI loci detected by generation 150 (Figure 5B). With such weak selection, less than ∼20% of true DMI loci (∼ 4 of 20) are detected within the mapped regions at generation 10, reaching a maximum of 50% (10 of 20) by generation 500 and with only ∼25% lying within estimated DMI regions even after this long timeframe (Figure 5B).

## Discussion

The genetic mapping of loci responsible for causing Dobzhansky-Muller incompatibilities (DMIs) in interspecies hybrids represents an ongoing challenge (Presgraves 2010b, Maheshwari and Barbash 2011). The value of natural hybrid populations toward this goal, as in taxa like house mice and swordtail fish (Powell et al. 2020, Forejt et al. 2021, Moran et al. 2021), raises the possibility that experimental interspecies hybrid populations derived from experimental evolution might provide an underexploited technique to complement more traditional genetic mapping approaches (Gray and Cutter 2014, Matute et al. 2020). We therefore simulated experimental evolution of hybrid populations to test for the power to detect DMI loci to inform the potential utility of an evolve-and-resequence experimental paradigm for this purpose. Overall, our observations indicate that mapping of DMIs with acceptable precision and accuracy using replicate hybrid populations is feasible, provided that the empirical system is capable of very rapid turnover of generations and that DMI fitness effects are neither too strong nor too weak. We found that the accuracy and precision of mapping trade off in our simulations of one-way incompatibilities, such that an intermediate duration of experimental evolution is preferred for such circumstances. In contrast, mapping of two-way incompatibilities benefits from extension of the duration of experimental evolution for as long as possible. As expected, DMI loci in regions of high recombination always are more amenable to efficient mapping. We also observed that little benefit accrued for DMI detection with greater population replication and that dominance interaction type exerted a strong influence just for two-way incompatibility detection. While an evolve-and-resequence approach to mapping DMI loci requires study systems with distinctive properties, with *Caenorhabditis* nematodes perhaps being especially suited to this technique (Gray and Cutter 2014, Castillo et al. 2015, Teotónio et al. 2017), our findings support a place for experimental evolution as an additional tool in the toolkit of speciation genetics researchers (White et al. 2020).

### Weak-to-moderate effect DMIs are more feasible to map for one-way incompatibilities

We find that strength of selection, duration of experimental evolution, and recombination rate all affect the power to localize one-way DMI loci and show a trade-off between precision and accuracy that is especially sensitive to stronger fitness effects (Figure 2). Our results indicate that if selection in hybrids arises from just a small number of strong-effect DMIs, then mapping with an evolve-and-resequence experiment would require propagation for an infeasible number of generations in most common study systems, albeit possible for organisms like *Caenorhabditis* nematodes (i.e. the best accuracy-precision trade-off of ∼100 generations would take ∼400 days). DMIs linked to high recombination regions, of course, can be mapped with greater power for a given duration of experimental evolution (Blanckaert and Bank 2018). A DMI locus pair that exerts weak to moderate effects, however, show more promise in yielding accurate and precise mapping resolution within 20 to 40 generations, with simulations showing the best trade-off at approximately 60 generations.

The higher precision of map estimates for weak and moderate-effect DMIs may be explained by the average time of DMI resolution: it takes longer to fix alleles that contribute to DMIs with weaker fitness effects (vertical dashed line in Figure S3). This slower resolution of DMI allows for more rounds of recombination to break up linked haplotype blocks surrounding the incompatibility loci. While this leads to estimates with higher precision, the resulting presence of fewer linked loci to draw information from compromises accuracy, demonstrating why the accuracies among weak selection coefficients do not exceed 90% in our simulations.

We did not find dominance relations between DMI loci to strongly affect the power to localize one-way DMI loci. In our design of an F_1_ hybrid starting population, selection can quickly “see” both dominant and recessive incompatibilities starting in the F_2_ generation. We expect dominance relations between DMI loci would exert a greater effect, however, when incompatibility alleles are not all present initially as heterozygotes. Incompatibility alleles starting at lower initial frequencies and in linkage disequilibrium might more closely match conditions simulated in other studies that report a stronger influence of DMI dominance interaction type (Turner and Harr 2014, Blanckaert and Bank 2018).

### Mapping of two-way incompatibilities benefits from long duration of experimental evolution

Two-way DMIs might be more common for species with greater overall divergence or for loci that experience rapid coevolution. In contrast to locus pairs involved in one-way incompatibilities, we observed both accuracy and precision of mapping two-way DMI loci to increase with the duration of experimental evolution (Figure 3; Figure 4). In two-way incompatibilities, all four alleles contribute to DMIs at the two loci and so selection affects both loci until both loci fix a single allele each. This requirement for fitness resolution in hybrid populations contrasts with one-way DMIs, which require fixation of only one locus, allowing polymorphic alleles at the partner DMI locus to then drift (Figure S3). The persistence of selection in the two-way scenario following fixation at one locus drives extreme allele frequencies at linked loci for both DMI loci to confer increased mapping power in later generations. Schumer et al. (Schumer et al. 2015) demonstrated a similar effect for what they referred to as symmetrical/coevolved incompatibilities (two-way) versus asymmetrical/neutral incompatibilities (one-way) in their simulations of hybrid speciation.

Mapping of DMI loci that cause two-way incompatibilities also differed from one-way incompatibilities in that we observed a strong influence of dominance. We observed high precision and accuracy of mapping for dominant two-way DMIs with moderate-to-small fitness effects in high recombination regions within a modest duration of experimental evolution (i.e., 20 to 40 generations, which could take ∼40 to ∼80 days in a real-world scenario applied to *Caenorhabditis* nematodes). Dominant two-way DMIs with the strongest fitness effects, however, suffer from poor precision until at least generation 150, even with high recombination. Recessive two-way DMIs, by contrast, show best mapping power when selection is strong-to-moderate. For example, strong-effect recessive DMIs have both precision and accuracy values above 80% by generation 20 for high recombination regions. These results indicate that mapping of two-way DMI loci may be most successful, for a given duration of experimental evolution, when the DMIs exert moderate fitness effects and lie in high recombination regions.

### Abundant one-way DMI pairs can generate substantial false positive mappings

While our analysis of single pairs of DMI loci proved instructive to assessing mapping power with an experimental evolution approach, the genomes of species are likely to encode many DMI loci (Masly and Presgraves 2007, Schumer et al. 2014). Consequently, we assessed DMI mapping in the context of hybrid simulations involving 10 pairs of one-way incompatibilities. Reflecting the accuracy-precision trade-off that we documented for pairs of one-way incompatibilities, we observed that longer durations of experimental evolution did not enhance the incidence of true positive mappings in the face of many DMI pairs in the simulated genomes. Allelic drift following fixation of one locus in a given DMI pair likely contributes to the inability of longer durations of experimental evolution to improve mapping confidence with many one-way DMI locus pairs encoded in genomes. The median width of the region surrounding estimated DMI locations narrows with longer durations of experimental evolution and smaller selection coefficients. However, the number of estimated DMIs increases over time beyond the true number of DMI loci, leading the fraction of true positives among estimated mapping locations to decline with the number of elapsed generations.

When selection on DMIs is strong, this trade-off between accuracy and precision leads to the best compromise near generation 100, and essentially all true DMI loci are detectable. Small-to-moderate fitness effects of the DMIs, however, lead to many more false positives in this timeframe and fewer true DMIs amenable to detection (i.e. lower accuracy), although the size of mapped regions are reasonably precise when they do overlap true DMI loci (mapping spans of 0.2% of chromosome length or less). These patterns indicate that if selection against many DMIs is suspected to be strong in real-world hybrid populations, then accurate and precise estimates of them could be achieved in a realistic amount of time in hybrid populations of organisms with rapid generation times. When selection against DMIs in hybrid genomes with many DMI pairs is weak-to-moderate, mapping also can be feasible if one is willing to accept the presence of many false positives. However, it is important to note that if selection against many DMIs is too strong, it could lead to population extinction. Population size was kept constant in these simulations and thus the potential for population extinction was not accounted for in these simulations, but could lead to lower power to detect DMIs in real-world settings. Moreover, hybrid genomes with high density of DMIs may lead to evolutionary reversion to a genomic state indistinguishable from just one of the parental species, as has been observed in experimental hybrid populations of *Drosophila* (Matute et al. 2020). This outcome could be exacerbated by higher genetic load in one of the parental species or by asymmetric potential for parental genotypes to be adaptive under lab conditions. Consequently, the power analyses of our simulations may overestimate mapping ability in the face of such additional real-world complications.

Finally, we found that the number of replicate populations exerted only a modest effect on the power to localize DMI loci in most cases with our approach. Consequently, excessive effort devoted to additional population replicates in real-world experiments often may not be worth-while. This observation contrasts with the positive influence of replicates in traditional quantitative trait locus mapping in evolve-and-resequence experiments (Kofler and Schlötterer 2014), and may reflect differences due to starting allele frequencies, additive versus epistatic selection, and alternative mapping inference techniques.

### Extensions to the investigation of DMIs with E&R

The populations modelled here simplify real-world DMIs with a number of potentially valuable extensions amenable to future work. For example, we only considered pairwise DMI interactions, but DMIs involving multiple interacting loci may be important in reproductive isolation (Dzur_JGejdosova et al. 2012, Schumer et al. 2014, Turner and Harr 2014, Tiago et al. 2015, Satokangas et al. 2020). Moreover, it remains unresolved how often species ought to exhibit one-way versus two-way versus multi-way incompatibilities, though this may depend in part on the amount of accumulated divergence. Future work also could also account for linkage between DMI loci, which is more likely when multiple genes are involved, rather than the encoding of partner loci on different chromosomes as in our simulations, or apply a co-ancestry disequilibrium scan approach to DMI detection in evolve and resequence studies (Schumer and Brandvain 2016). Related to the issue of linkage is genome collinearity between species, such that the genomes of real species may show inversions and other rearrangements (Faria and Navarro 2010, Fuller et al. 2018) that could compromise the power to localize DMIs relative to simulations like ours that presume perfect synteny.

Another important extension of this work would be sex-linkage or sex-limited expression of DMI loci (Coyne and Orr 1998b). Autosomal DMI loci with sex-limited expression will expose their fitness effects in only half of a population, so the power to detect them may resemble the small-effect DMIs simulated here. There is no real consensus, however, on what the fitness effects of DMI locus pairs should be, and thus it also will prove valuable to explore a broader spectrum of selection coefficients. Moreover, the DMI alleles that we modelled are neutral in their respective parental species’ backgrounds, but adaptive alleles could also be modelled. Schumer et al. (Schumer et al. 2015) note that it is currently not known whether adaptive or neutral DMI alleles are more prevalent, and they modelled hybrid populations with both types of incompatibilities. In the adaptive case, alleles at distinct DMI loci are more likely to fix for the same parental species’ allele combination (Schumer et al. 2015). Consequently, the outcomes of our two-way DMIs that also tend to fix DMI alleles in this way may capture some of the features of adaptive DMI allele introgression.

Another potential extension of this work would be to investigate whether the results could apply to detecting other forms of epistasis, as may be important among loci within species (Whitlock et al. 1995, Montooth et al. 2010). Detecting parallel allele frequency changes across replicate populations under specific parameter spaces might permit detection of loci implicated in meiotic drive and other intragenomic conflicts (Kofler et al. 2012, Rice 2013).

## Conclusion

The ability to accurately and precisely detect DMI loci presents a challenge to characterizing the genetic underpinnings of reproductive isolation between species. Our exploration of the feasibility of experimental evolution with an evolve-and-resequence approach to map DMIs found that it will be most successful for modest fitness effects of DMIs caused by one-way incompatibilities, rather than two-way incompatibilities, that reside in high recombination regions. DMI mapping in genomes that contain many locus pairs are subject to a trade-off between precision and accuracy that may be acceptable for strong-selection coefficients or if one is willing to accept the presence of many false positives. Unlike the effects of differing selection coefficients and duration of experimental evolution, the number of replicate populations exerted only a modest effect on the power to localize DMI loci in most cases with our approach. Altogether, these findings demonstrate the plausibility of DMI mapping over feasible durations of experimental evolution with hybrid populations of organisms that have rapid turnover of generations, such as *Caenorhabditis* nematodes, to provide an addition to the toolkit of speciation genetics researchers.

## Acknowledgments

We thank Santiago Sánchez-Ramírez and Ben Haller for software consultation. Computations were performed on the Niagara supercomputer at the SciNet HPC Consortium (funded by the Canada Foundation for Innovation, the Government of Ontario, Ontario Research Fund - Research Excellence, and the University of Toronto). A.D.C. is supported by a Discovery Grant from the Natural Sciences and Engineering Research Council of Canada.

## Data accessibility and benefit-sharing

Data accessibility statement: Simulation and analysis scripts are available at github.com/Cutterlab/Generating-DMI-Estimates Benefit-sharing statement: N/A

## Author contributions

N.S. designed and performed the research, analyzed data, and wrote the paper. A.D.C. designed the research and wrote the paper.

## Supplementary

**Figure S1.**
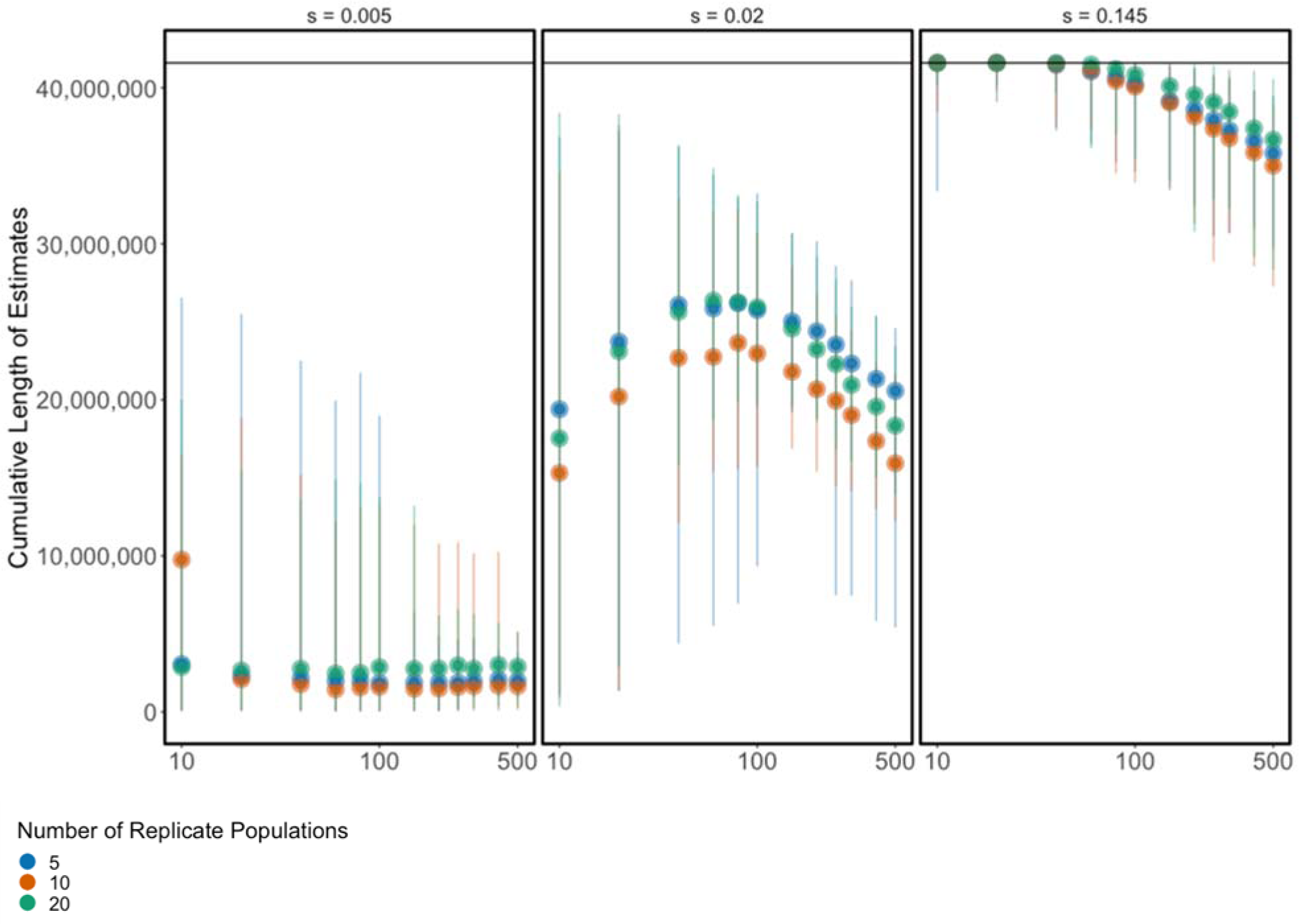
Cumulative length of mapped DMI estimates as a function of the duration of experimental evolution (log scale generations) for 20 DMI loci that express one-way pairwise incompatibilities, inferred from 5, 10, and 20 replicate populations. Error bars correspond to 2.5^th^ to 97.5^th^ quantiles.

**Figure S2.**
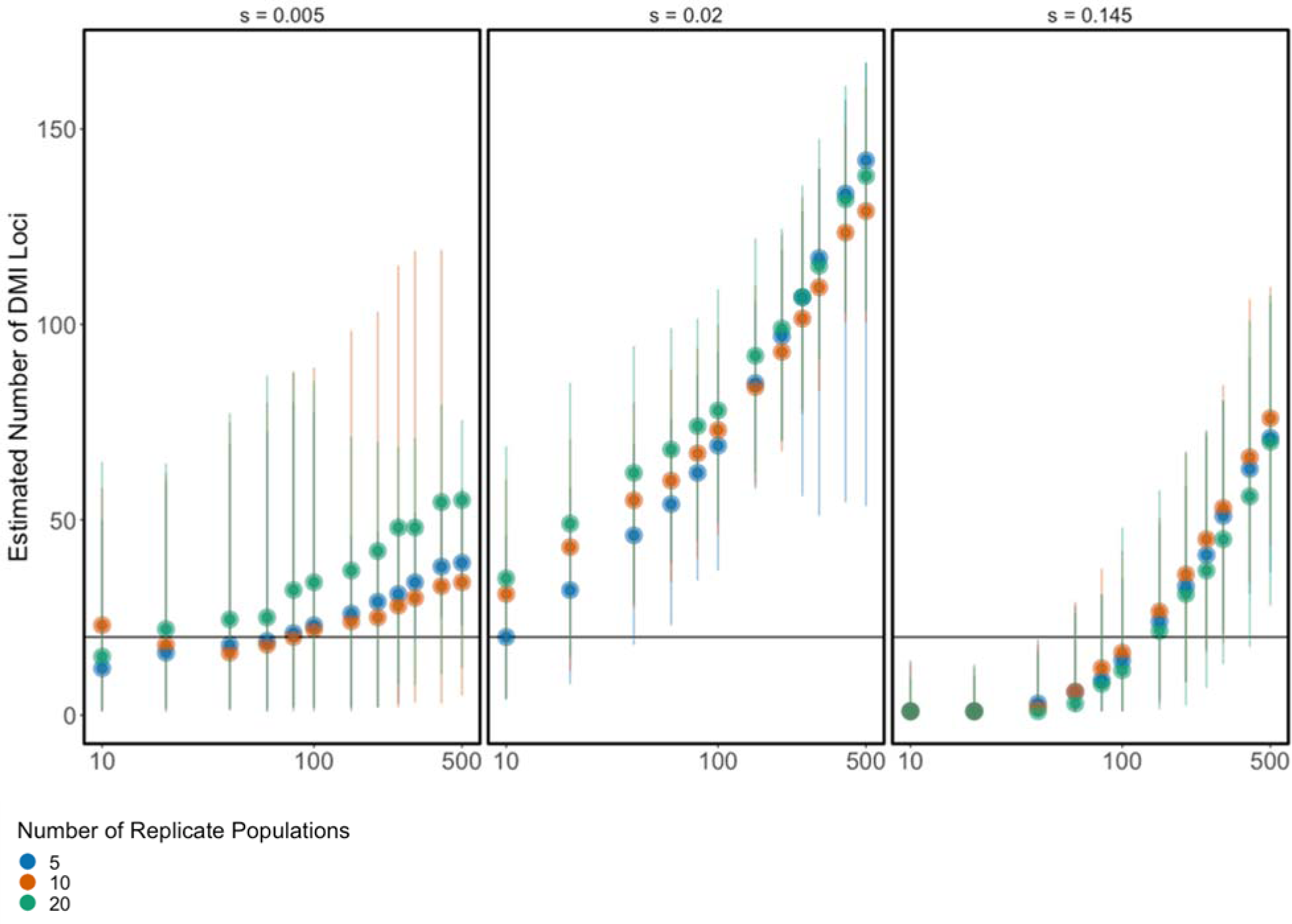
Estimated number of DMI loci as a function of the duration of experimental evolution (log scale generations) for 20 DMI loci that express pairwise one-way incompatibilities, inferred from 5, 10, and 20 replicate populations. Error bars correspond to 2.5^th^ to 97.5^th^ quantiles.

**Figure S3.**
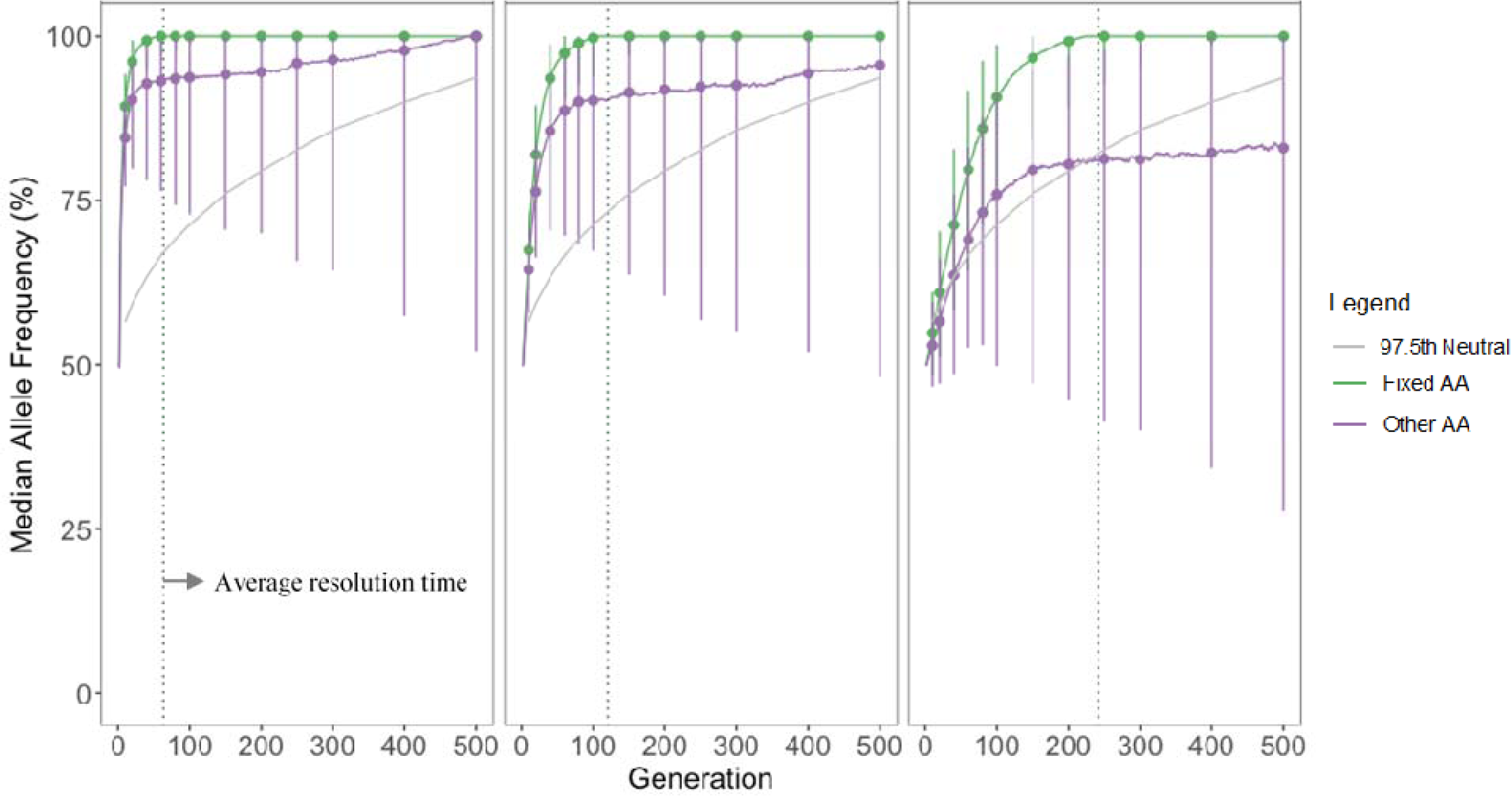
Median allele frequencies at each locus in a DMI pair for one-way incompatibilities. The ancestral allele (AA) that fixed first in each population (green line) causes DMI resolution. The frequency of the other ancestral allele at the other locus (purple line) are then subject only to genetic drift in subsequent generations. Frequency data were collected at each generation 1 to 500, collected separately from the simulation runs used to generate estimates, but resembled them in every way except for the exclusion of the 20,000 neutral loci. In each population, the fixed ancestral allele can either be from parental species 1 or 2. Median values are calculated from 1000 replicate populations that contain a one-way dominant DMI with a selection coefficient of 0.05, 0.2, or 0.8. The error bars for 12 timepoints are the frequency 95% interquantiles ranging from the 2.5^th^ and 97.5^th^ quantiles. The dashed vertical line corresponds to the average timepoint at which DMI resolution occurs across replicates. The gray curve indicates the 97.5^th^ quantiles of allele frequencies from 1000 populations lacking fitness-affecting DMI loci.

## References

Barton, N. H., and G. M. Hewitt. 1989. Adaptation, speciation and hybrid zones. Nature 341:497–503.

Blackman, B. K. 2016. Speciation genes. Pages 166-175 in R. M. Kliman, editor. Encyclopedia of Evolutionary Biology. Academic Press, Oxford.

Blanckaert, A., and C. Bank. 2018. In search of the Goldilocks zone for hybrid speciation. PLoS Genetics 14:e1007613.

Bundus, J. D., D. Wang, and A. D. Cutter. 2018. Genetic basis to hybrid inviability is more complex than hybrid male sterility in *Caenorhabditis* nematodes. Heredity 121:169–181.

Castillo, D. M., and D. A. Barbash. 2017. Moving speciation genetics forward: modern techniques build on foundational studies in *Drosophila*. Genetics 207:825–842.

Castillo, D. M., M. K. Burger, C. M. Lively, and L. F. Delph. 2015. Experimental evolution: Assortative mating and sexual selection, independent of local adaptation, lead to reproductive isolation in the nematode *Caenorhabditis remanei*. Evolution 69:3141–3155.

Coughlan, J. M., and D. R. Matute. 2020. The importance of intrinsic postzygotic barriers throughout the speciation process. Philosophical Transactions of the Royal Society B: Biological Sciences 375:20190533.

Coyne, J. A., and H. A. Orr. 1998a. The evolutionary genetics of speciation. Philosophical Transactions of the Royal Society of London B: Biological Sciences 353:287–305.

Coyne, J. A., and H. A. Orr. 1998b. The evolutionary genetics of speciation. Philosophical Transactions of the Royal Society of London Series B-Biological Sciences 353:287–305.

Cutter, A. D. 2018. X exceptionalism in *Caenorhabditis* speciation. Molecular Ecology 27:3925–3934.

De La Torre, A., P. K. Ingvarsson, and S. N. Aitken. 2015. Genetic architecture and genomic patterns of gene flow between hybridizing species of *Picea*. Heredity 115:153–164.

Dey, A., Q. Jin, Y.-C. Chen, and A. D. Cutter. 2014. Gonad morphogenesis defects drive hybrid male sterility in asymmetric hybrid breakdown of *Caenorhabditis* nematodes. Evolution and Development 16:362–372.

Dzur_JGejdosova, M., P. Simecek, S. Gregorova, T. Bhattacharyya, and J. Forejt. 2012. Dissecting the genetic architecture of F1 hybrid sterility in house mice. Evolution 66:3321–3335.

Faria, R., and A. Navarro. 2010. Chromosomal speciation revisited: rearranging theory with pieces of evidence. Trends in Ecology & Evolution 25:660–669.

Ferretti, L., S. E. Ramos-Onsins, and M. Pérez-Enciso. 2013. Population genomics from pool sequencing. Molecular Ecology 22:5561–5576.

Fitzpatrick, B. M. 2013. Alternative forms for genomic clines. Ecology and Evolution 3:1951–1966.

Forejt, J., P. Jansa, and E. Parvanov. 2021. Hybrid sterility genes in mice (Mus musculus): a peculiar case of PRDM9 incompatibility. Trends in Genetics 37:1095–1108.

Fuller, Z. L., C. J. Leonard, R. E. Young, S. W. Schaeffer, and N. Phadnis. 2018. Ancestral polymorphisms explain the role of chromosomal inversions in speciation. PLoS Genetics 14:e1007526.

Gompert, Z., and C. A. Buerkle. 2011. Bayesian estimation of genomic clines. Molecular Ecology 20:2111–2127.

Gray, J. C., and A. D. Cutter. 2014. Mainstreaming *Caenorhabditis elegans* in experimental evolution. Proceedings of the Royal Society of London B: Biological Sciences 281:20133055.

Haller, B. C., and P. W. Messer. 2019. SLiM 3: Forward genetic simulations beyond the Wright– Fisher model. Molecular Biology and Evolution 36:632–637.

Kofler, R., A. J. Betancourt, and C. Schlötterer. 2012. Sequencing of pooled DNA samples (Pool-Seq) uncovers complex dynamics of transposable element insertions in *Drosophila melanogaster*. PLoS Genetics 8:e1002487.

Kofler, R., R. V. Pandey, and C. Schlotterer. 2011. PoPoolation2: identifying differentiation between populations using sequencing of pooled DNA samples (Pool-Seq). Bioinformatics 27:3435–3436.

Kofler, R., and C. Schlötterer. 2014. A guide for the design of evolve and resequencing studies. Molecular Biology and Evolution 31:474–483.

Li, C., Z. Wang, and J. Zhang. 2013. Toward genome-wide identification of Bateson– Dobzhansky–Muller incompatibilities in yeast: a simulation study. Genome Biology and Evolution 5:1261–1272.

Long, A., G. Liti, A. Luptak, and O. Tenaillon. 2015. Elucidating the molecular architecture of adaptation via evolve and resequence experiments. Nature Reviews Genetics 16:567–582.

Maharjan, R. P., and T. Ferenci. 2013. Epistatic interactions determine the mutational pathways and coexistence of lineages in clonal *Escherichia coli* populations. Evolution 67:2762–2768.

Maheshwari, S., and D. A. Barbash. 2011. The genetics of hybrid incompatibilities. Annual Review of Genetics 45:331–355.

Masly, J. P., and D. C. Presgraves. 2007. High-resolution genome-wide dissection of the two rules of speciation in *Drosophila*. PLoS Biology 5:e243.

Matute, D. R., A. A. Comeault, E. Earley, A. Serrato-Capuchina, D. Peede, A. Monroy-Eklund, W. Huang, C. D. Jones, T. F. C. Mackay, and J. A. Coyne. 2020. Rapid and predictable evolution of admixed populations between two *Drosophila* species pairs. Genetics 214:211.

Montooth, K. L., C. D. Meiklejohn, D. N. Abt, and D. M. Rand. 2010. Mitochondrial-nuclear epistasis affects fitness within species but does not contribute to fixed incompatibilities between specis of Drosophila. Evolution 64:3364–3379.

Moran, B. M., C. Payne, Q. Langdon, D. L. Powell, Y. Brandvain, and M. Schumer. 2021. The genomic consequences of hybridization. eLife 10:e69016.

Payseur, B. A. 2010. Using differential introgression in hybrid zones to identify genomic regions involved in speciation. Molecular Ecology Resources 10:806–820.

Pereira, R. J., T. G. Lima, N. T. Pierce-Ward, L. Chao, and R. S. Burton. 2021. Recovery from hybrid breakdown reveals a complex genetic architecture of mitonuclear incompatibilities. Molecular Ecology 30:6403–6416.

Ponce, M., R. v. Zon, S. Northrup, D. Gruner, J. Chen, F. Ertinaz, A. Fedoseev, L. Groer, F. Mao, B. C. Mundim, M. Nolta, J. Pinto, M. Saldarriaga, V. Slavnic, E. Spence, C.-H. Yu, and W. R. Peltier. 2019. Deploying a Top-100 supercomputer for large parallel workloads: the Niagara Supercomputer. Page Article 34 Proceedings of the Practice and Experience in Advanced Research Computing on Rise of the Machines (learning). Association for Computing Machinery, Chicago, IL, USA.

Powell, D. L., M. García-Olazábal, M. Keegan, P. Reilly, K. Du, A. P. Díaz-Loyo, S. Banerjee, D. Blakkan, D. Reich, P. Andolfatto, G. G. Rosenthal, M. Schartl, and M. Schumer. 2020. Natural hybridization reveals incompatible alleles that cause melanoma in swordtail fish. Science 368:731–736.

Presgraves, D. C. 2010a. Darwin and the origin of interspecific genetic incompatibilities. American Naturalist 176 Suppl 1:S45–S60.

Presgraves, D. C. 2010b. The molecular evolutionary basis of species formation. Nature Reviews Genetics 11:175–180.

Rice, W. R. 2013. Nothing in genetics makes sense except in light of genomic conflict. Annual Review of Ecology, Evolution, and Systematics 44:217–237.

Rockman, M. V., and L. Kruglyak. 2009. Recombinational landscape and population genomics of *C. elegans*. PLoS Genetics 5:e1000419.

Satokangas, I., S. H. Martin, H. Helanterä, J. Saramäki, and J. Kulmuni. 2020. Multi-locus interactions and the build-up of reproductive isolation. Philosophical Transactions of the Royal Society B: Biological Sciences 375:20190543.

Savolainen, O., M. Lascoux, and J. Merilä. 2013. Ecological genomics of local adaptation. Nature Reviews Genetics 14:807–820.

Schumer, M., and Y. Brandvain. 2016. Determining epistatic selection in admixed populations. Molecular Ecology 25:2577–2591.

Schumer, M., R. Cui, D. L. Powell, R. Dresner, G. G. Rosenthal, and P. Andolfatto. 2014. High-resolution mapping reveals hundreds of genetic incompatibilities in hybridizing fish species. eLife 3:e02535.

Schumer, M., R. Cui, G. G. Rosenthal, and P. Andolfatto. 2015. Reproductive isolation of hybrid populations driven by genetic incompatibilities. PLoS Genetics 11:e1005041.

Schumer, M., D. L. Powell, and R. Corbett-Detig. 2020. Versatile simulations of admixture and accurate local ancestry inference with mixnmatch and ancestryinfer. Molecular Ecology Resources 20:1141–1151.

Stankowski, S., and M. Ravinet. 2021. Defining the speciation continuum. Evolution 75:1256–1273.

Teotónio, H., S. Estes, P. C. Phillips, and C. F. Baer. 2017. Experimental evolution with *Caenorhabditis* nematodes. Genetics 206:691.

Teterina, A. A., J. H. Willis, and P. C. Phillips. 2020. Chromosome-level assembly of the *Caenorhabditis remanei* genome reveals conserved patterns of nematode genome organization. Genetics 214:769–780.

Tiago, P., E. B. Kevin, and B. R. A. Ricardo. 2015. Emergent speciation by multiple Dobzhansky–Muller Incompatibilities. bioRxiv:008268.

Turner, L. M., and B. Harr. 2014. Genome-wide mapping in a house mouse hybrid zone reveals hybrid sterility loci and Dobzhansky-Muller interactions. eLife 3:e02504.

Turner, T. L., E. C. Bourne, E. J. Von Wettberg, T. T. Hu, and S. V. Nuzhdin. 2010. Population resequencing reveals local adaptation of *Arabidopsis lyrata* to serpentine soils. Nature Genetics **In press**.

Turner, T. L., A. D. Stewart, A. T. Fields, W. R. Rice, and A. M. Tarone. 2011. Population-based resequencing of experimentally evolved populations reveals the genetic basis of body size variation in *Drosophila melanogaster*. PLoS Genetics 7:e1001336.

White, N. J., R. R. Snook, and I. Eyres. 2020. The past and future of experimental speciation Trends in Ecology & Evolution 35:10–21.

Whitlock, M. C., P. C. Phillips, F. B. G. Moore, and S. J. Tonsor. 1995. Multiple fitness peaks and epistasis. Annual Review of Ecology and Systematics 26:601–629.

Woodruff, G. C., O. Eke, S. E. Baird, M. A. Felix, and E. S. Haag. 2010. Insights into species divergence and the evolution of hermaphroditism from fertile interspecies hybrids of *Caenorhabditis* nematodes. Genetics 186:997–1012.

